# Inferring Phylogenetic Trees of Cancer Evolution from Longitudinal Single-Cell Copy Number Profiles

**DOI:** 10.1101/2025.03.17.643834

**Authors:** Yushu Liu, Luay Nakhleh

## Abstract

Understanding evolutionary dynamics is critical for unraveling the complex progression of diseases such as cancer. Cancer evolution is inherently a temporal process driven by the accumulation of mutations and clonal expansions over time. Traditional phylogenetic methods often rely solely on static, cross-sectional data, limiting their ability to infer the timing of key evolutionary events. To address this challenge, we developed NestedBD-Long, a novel method that integrates temporal data from longitudinal sampling into phylogenetic analyses using the birthdeath evolutionary model on copy numbers. This approach allows for the direct mapping of real-world time onto inferred evolutionary trees, providing a clearer and more accurate representation of cancer’s evolutionary trajectory. Evaluations demonstrate that NestedBD-Long outperforms traditional approaches, with accuracy improving as the number of temporal sampling points increases. This advancement provides a powerful framework for studying tumor progression, treatment resistance, and metastatic spread by capturing the dynamics between evolutionary events and real-world timelines. NestedBD-Long is available at https://github.com/Androstane/NestedBD.

## 1 Introduction

Copy number alterations (CNAs), involving amplifications or deletions of DNA segments, are key genomic events driving cancer progression [6]. These alterations can disrupt gene dosage, leading to either the overexpression or loss of critical genes, ultimately contributing to uncontrolled cell growth and tumor evolution [3,19]. The study of CNAs is crucial for understanding the mechanisms underlying tumorigenesis and identifying potential therapeutic targets in cancer [8].

Single-cell DNA sequencing (scDNAseq) has revolutionized cancer genomics by enabling the study of genomic heterogeneity at the resolution of individual cells [27,11]. This technology provides a detailed view of clonal architecture, capturing the diversity of copy number alterations (CNAs) within tumors and facilitating the reconstruction of their evolutionary trajectories. scDNAseq has been applied to various cancer types, yielding valuable insights into tumor progression and therapy resistance [17,9].

In recent years, researchers have increasingly employed longitudinal singlecell sequencing—where sequencing is performed at multiple time points—to monitor temporal changes in genomic features throughout disease progression. This approach offers a dynamic perspective on tumor evolution, particularly under the selective pressures of treatment [23,30]. By analyzing genomic alterations before, during, and after therapeutic interventions, researchers can track the emergence of resistant subclones, identify critical evolutionary bottlenecks, and refine treatment strategies. Longitudinal sequencing enables the identification of key evolutionary events, such as the rise of therapy-resistant subclones, thus providing opportunities for more precise therapeutic adjustments.

Despite its promise, there remain limited computational methods that fully leverage longitudinal sequencing data in phylogenetic inference. For example, while tools such as LACE and scLongTree [20,1,15] can infer phylogenetic trees from longitudinal single-nucleotide variant (SNV) data, they assume uniform time intervals between consecutive sampling points—failing to account for cases where one interval is significantly longer than another. Moreover, to our knowledge, no existing methodologies are explicitly designed for the inference of longitudinal single-cell CNA data.

In this paper, we address this gap by introducing a novel approach, NestedBD-Long, for inferring phylogenetic trees from longitudinal single-cell CNA data, explicitly incorporating real-world temporal information (e.g., days or months) via a birth-death evolutionary model introduced in [18]. NestedBD-Long is based on NestedBD, utilizing the same evolutionary model and leveraging BEAST2 [2] for Bayesian inference, but extends its capability to handle longitudinal data. This feature enables a time-resolved analysis of tumor evolution, distinguishing our method from existing tools such as MEDICC2 [14] and Lazac [22], which can process CNA data but do not integrate explicit temporal sampling. Figure 1 provides an overview of our proposed method.

**Fig. 1.**
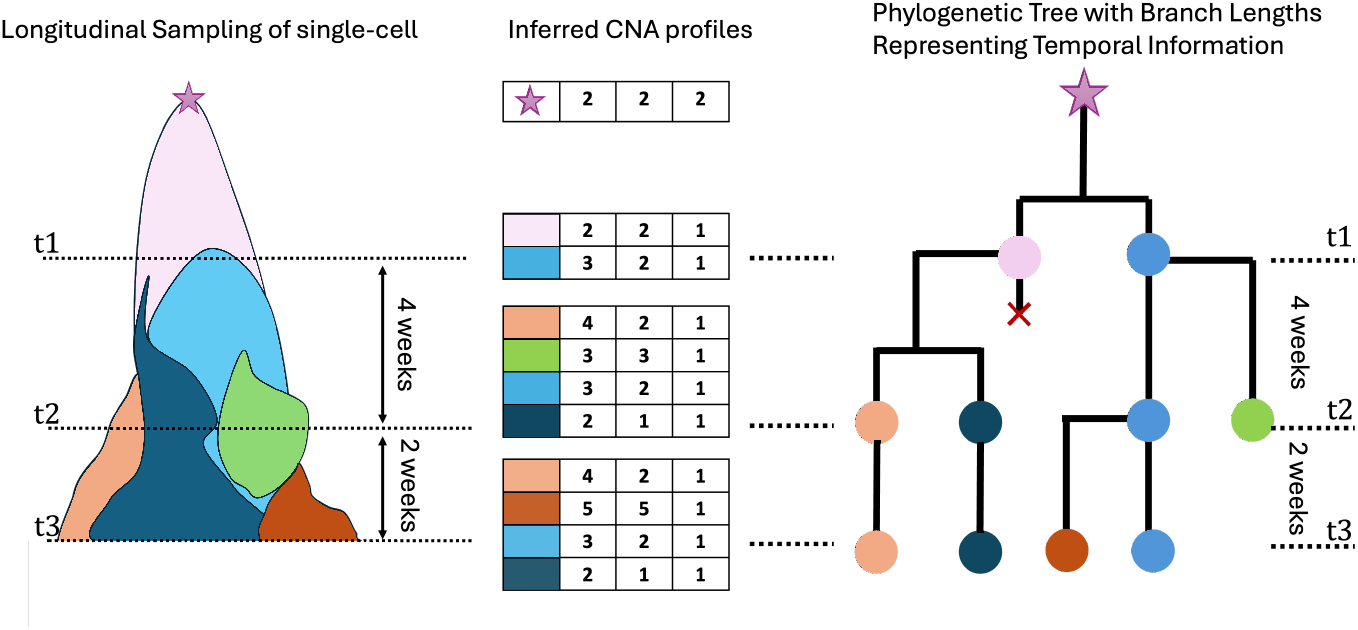
Overview of NestedBD-Long. From binned copy number profiles collected at multiple time points, the method infers a single-cell phylogenetic tree with branch lengths mapped to real-world time and branch-specific mutation rates.

We evaluated NestedBD-Long on both simulated and real biological datasets. The simulations were designed to replicate the complexities of tumor evolution across multiple time points, capturing key genomic alterations and clonal dynamics. Real datasets provided a benchmark for assessing the practical applicability of our approach in real-world cancer studies. Our results demonstrate that explicitly incorporating temporal information substantially improves phylogenetic accuracy, particularly as the number of temporal sampling points increases. Moreover, NestedBD-Long consistently outperforms traditional approaches by more accurately reconstructing evolutionary trajectories and providing deeper insights into tumor progression. This capability to map inferred evolutionary time onto real-world temporal scales represents a significant advancement in cancer phylogenetics, facilitating more precise analyses of tumor evolution, treatment response, and resistance mechanisms.

NestedBD-Long is available at https://github.com/Androstane/NestedBD.

## 2 Methods

### 2.1 Temporal Integration of Longitudinal CNA Data

To infer phylogenetic trees from longitudinal CNA data, we employed a birthdeath skyline model [10] that incorporates sampled ancestors. While this model was employed in the context of epidemiology and fossil calibration, we adapt it here given the analogy between ancestral samples and fossil data. This model accounts for serially sampled cells from different time points and allows sampled lineages to persist and contribute to subsequent clonal expansions. The model is parameterized by four rates:

– the *birth rate* (*λ*), representing the probability of speciation;
– the *death rate* (*µ*), representing the probability of extinction;
– the *sampling rate* (*ψ*), representing the probability of observing a lineage through sampling; and,
– the *post-sampling removal probability* (*r*), representing the probability that a sampled lineage is removed from further evolution. Each sampled lineage is either removed with probability *r* or remains in the population, contributing to future diversification as a sampled ancestor.

This results in three classes of nodes in the reconstructed phylogeny *T* :

– bifurcation nodes, representing cell divisions;
– sampled tip nodes, corresponding to sampled lineages at the tips; and,
– sampled internal nodes, denoting lineages that were sampled but continued to evolve.

Given the model specification, the probability of observing a phylogenetic tree *T* can be computed as:

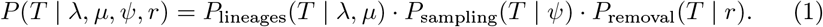

This equation decomposes the likelihood into three independent terms. The first term, *P*_lineages_(*T* |*λ, µ*), describes the probability of generating the observed tree topology given the birth and death rates. It accounts for the balance between clonal expansion and extinction, which determines the overall structure of the tree. A higher birth rate relative to the death rate results in larger trees with more bifurcations, while a high extinction rate leads to smaller trees with frequent lineage loss. The second term, *P*_sampling_(*T* |*ψ*), represents the likelihood of observing a lineage through sampling at different time points. Since tumor samples are collected longitudinally, this term conditions the inference on the observed data, incorporating the probability of detecting a given lineage at a particular time. The third term, *P*_removal_(*T* |*r*), accounts for whether sampled lineages persist in the tumor or are removed from further evolution. When *r* = 1, all sampled cells are assumed to be terminal, meaning that once a cell is sampled, it does not give rise to any new descendants in the reconstructed tree. When *r <* 1, some sampled cells persist and may be resampled at a later time point, leading to the presence of sampled ancestor nodes in the tree.

To infer phylogenies under this model, we employ a Bayesian inference framework implemented in BEAST2 [2], incorporating the evolutionary model from NestedBD [18]. Specifically, we estimate the posterior distribution of phylogenetic trees using a combination of the birth-death evolutionary model of copy number profiles and the birth-death skyline model leveraging Markov Chain Monte Carlo (MCMC) for parameter inference. The posterior distribution is given by:

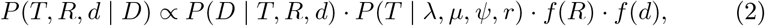

the extinction fraction where *D* is the estimated copy number profile, *d* is the distance between the common ancestor of all cells and its diploid ancestor, and *R* is the collection of parameters that define the clock model on *T* . We specified the sampled ancestor tree model using the Sampled Ancestor package in BEAST2 [10]. The diversification rate (*r*_d_ = *λ* − *µ*) was assigned a Uniform(0, 1000000) prior, while *(r*_e_ = *µ/λ*) and the sampling rate (*ψ*) were assigned Uniform(0, 1) priors. We assumed the post-sampling removal probability *r* = 1, consistent with the lineage removal process described in Section ‘Longitudinal Sampling’ and the biological constraint that a cell, once removed during sequencing, can no longer undergo further evolution. Additional parameter specifications follow those in NestedBD [18].

Given the likelihood, we employ standard tree moves in [2] to explore the tree space. The tree moves used in this study with their weights also follow specification in [18].

While the model allows for rate variation over time, we assume constant rates to mitigate potential parameter identifiability issues. For simulated datasets, node ages were defined by preset sampling times, while for biological datasets, tree heights were rescaled so that the oldest sampled cell was assigned an age of 1. A figure illustrating a sampled ancestor tree versus a reconstructed tree under the specified parameters is shown in Figure 2.

**Fig. 2.**
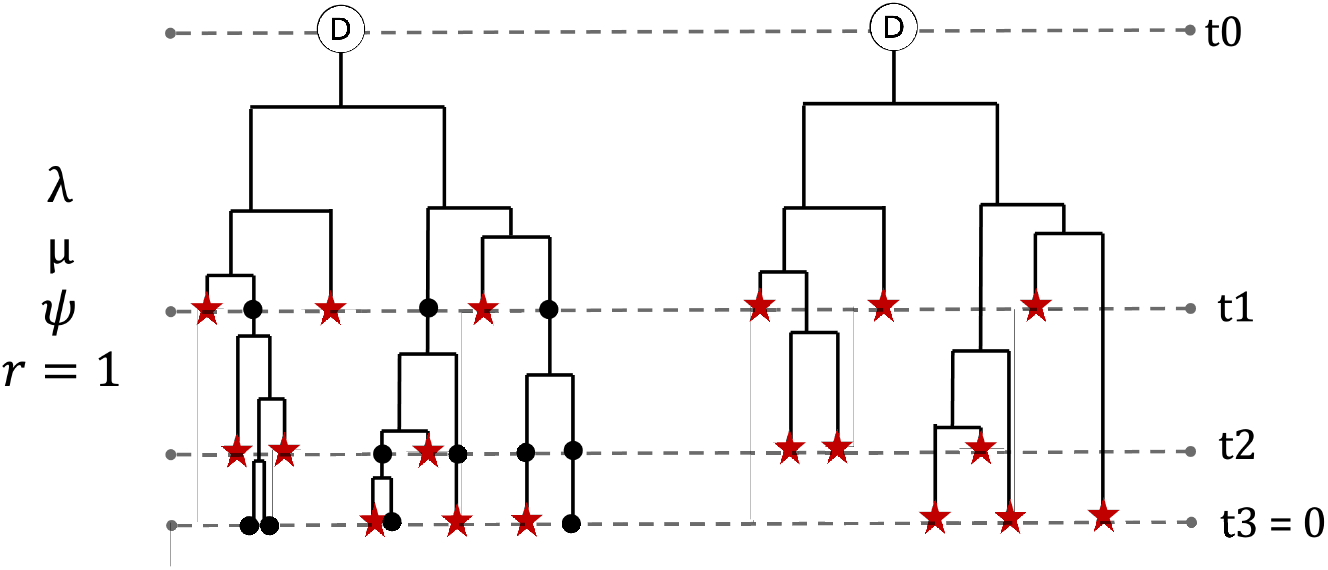
Full tree versus reconstructed tree. The left panel is a full tree generated by the sampled ancestor birth-death process with a post-sampling removal probability of *r* = 1, meaning all sampled lineages are terminal. The tree follows a birth-death process, where each lineage evolves with a birth rate *λ* and a death rate *µ* at each interval. At each sampling time point *t*_1_ to *t*_3_, all existing lineages are independently sampled with probability *ψ*. Red stars represent sampled nodes that do not contribute to future evolution (or sampled tips at *t*_3_), while black dots represent unsampled lineages that continued to evolve(or unsampled tips at *t*_3_). The right panel shows the corresponding reconstructed tree, which includes only observed sampled nodes and their ancestral relationships. The node labeled *D* at *t*_0_ represents the healthy diploid cell, serving as the ancestral state before clonal evolution.

### 2.2 Simulation

To evaluate the robustness of NestedBD-Long under diverse conditions, we varied simulation parameters, including temporal sampling intervals, evolutionary scenarios, and noise levels in the simulated CNA profiles. For each parameter combination, we generated 10 independent replicates to ensure statistical reliability. The performance of the methods is then assessed by aggregating results across these replicates, allowing for a comprehensive comparison of accuracy and consistency across different evolutionary scenarios.

### Simulating Phylogenetic Trees

As the first step, we generated an ultrametric tree with 1000 extant leaves under a birth-death model with equal birth and death rates using the simulator described in [25]. The root node was set as a diploid state without any CNAs, while each leaf node represented a singlecell sample from the patient. Internal nodes in the simulated trees represented ancestral cells that existed in the past but were not directly sampled. To maintain consistency across simulations, the branch lengths of the generated trees were scaled such that all trees had a total height of 1, ensuring that the expected number of evolutionary events remained uniform regardless of the number of sampled cells. To model potential sampling time points, we introduced additional internal nodes along each lineage at specific heights of 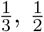 and 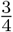 of the tree’s height, to represent potential sampling events.

#### Simulating CNAs

Given an input tree with 1000 leaves, we used the simulator described in [28] to model CNA events along the branches. The simulations were performed with *x* = 22 chromosomes, a mean of *λ* = 2 CNAs per edge, and a bin size of *b* = 500 kbp. The number of clones representing distinct subclonal lineages was set to *c* ∈{0, 4, 10}. Clonality was introduced by selecting ancestral nodes to define diverging subclonal populations, with selection based on subtree size or branch length. Boundary error rates, representing misidentifications at CNA breakpoints, were defined as *r*_b_ ∈ {0.08, 0.14} for low-noise and high-noise datasets, respectively. Additionally, a fixed jitter error rate of *r*_j_ = 0.15 was applied to introduce random fluctuations in copy number states, reflecting read count nonuniformity in scDNA-seq data.

#### Longitudinal Sampling

Cell sampling was performed from the simulated tree in Section ‘Simulating Phylogenetic Trees’ with 1000 leaves and three potential sampling times set at heights of 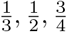 of the tree’s height. We varied the number of sampling points, *n* = 1, 2, 3, 4, ensuring that for each *n*, the current time *t*_0_ (corresponding to the leaves) was always included, along with *n* −1 additional time points randomly selected from the predefined heights. Sampling was conducted sequentially, from the earliest to the latest time points, by randomly selecting 30 nodes. To mirror biological sequencing, once a cell is selected, its entire lineage, including all descendant cells, is removed from the tree. This process was repeated at each time point until 30 cells were obtained for all designated sampling times. If an insufficient number of cells remained at a later time point due to earlier sampling, the procedure was restarted until a successful sampling process could be completed.

### 2.3 Evaluating Inference Accuracy

We assessed the topological accuracy of each method by calculating the normalized Robinson-Foulds (RF) distance [21] between the inferred and true topologies. For methods that provide a direct point estimate of the tree topology, we used the inferred tree as is. In cases where a method employs Bayesian inference and offers a posterior distribution rather than a point estimate, we summarized this distribution by taking 2,000 samples from the MCMC chain to compute the inferred topology and branch lengths. We then determined the maximum clade credibility tree (MCC), defined as the tree with the highest product of posterior clade probabilities, and summarized branch lengths using the median node heights across the samples.

## 3 Results

To our knowledge, no existing method explicitly infers phylogenetic trees from longitudinal single-cell CNA data. While LACE[1] and scLongtree[15] incorporate longitudinal information, they use SNVs rather than CNAs as input and are therefore not applicable to this study (though generating CNA and SNV data from the same sample and comparing the methods is a direction for future research). Consequently, we compare NestedBD-Long to MEDICC2[13], Lazac[22] and NestedBD[18], which infer phylogenetic trees from single-cell CNA data but do not explicitly model longitudinal sampling. We evaluate the performance of these methods on simulated datasets by assessing the accuracy of inferred trees under varying sampling time points, clonality levels, and CNA profiling error rates. Additionally, we apply these methods to biological single-cell copy number profiles from TNBC patients to compare their inferred phylogenies and examine how well they capture temporal structure.

### 3.1 Performance on simulated data

#### Accuracy with Varying Sampling Time Points

We evaluated the performance of each method on simulated copy number profiles by computing the normalized RF distance between the inferred and ground truth trees. This analysis assesses the impact of increasing the number of sampling time points on topological reconstruction accuracy. The results are summarized in Figure 3.

**Fig. 3.**
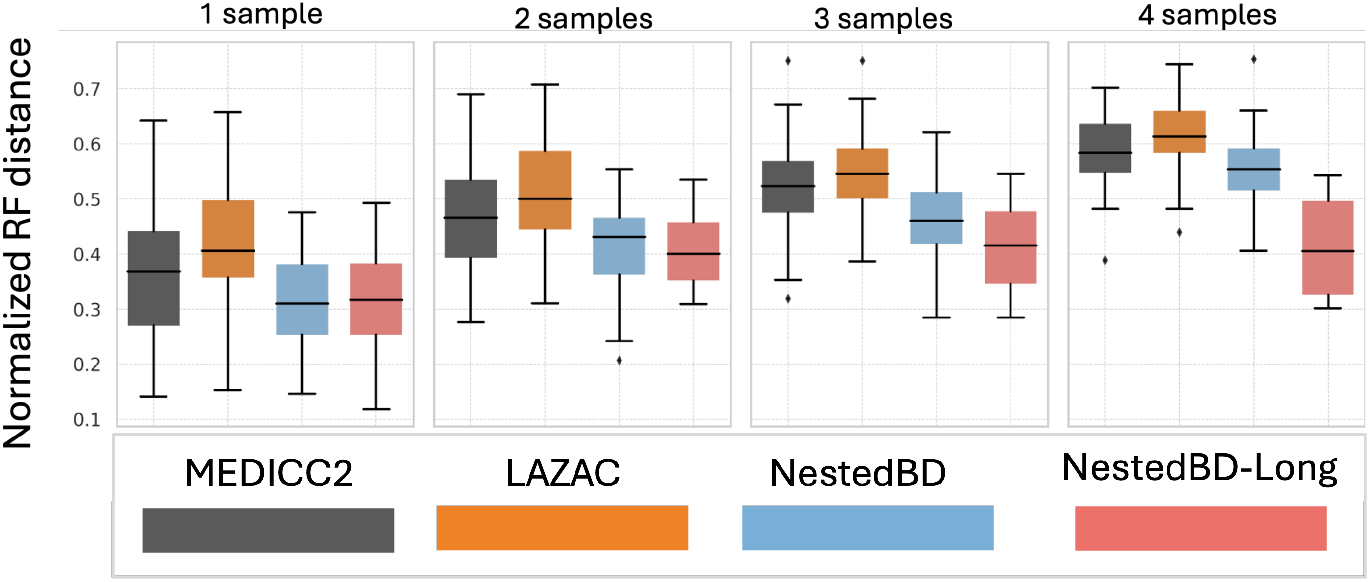
Accuracy of the inferred trees on the simulated data by number of sampling time points. The box plots summarize the distributions of normalized RF distances between the ground truth and inferred trees for each method using 10 replicates.

The results indicate that the Bayesian methods NestedBD and NestedBD-Long outperform the distance-based methods Lazac and MEDICC2. Notably, NestedBD-Long maintains consistent performance even as the number of longitudinal sampling points increases, underscoring the benefits of integrating temporal information into tree inference. Specifically, NestedBD-Long maintains a narrow range of normalized RF distances across various sampling points, whereas other methods experience wider variability and increased RF distances as the number of sampling points grows. The gap between methods opens up as more sampling points are introduced, highlighting the challenges non-Bayesian methods face in handling the added complexity. In summary, these findings demonstrate that leveraging temporal information through NestedBD-Long can improve the accuracy of phylogenetic analyses in longitudinal studies.

#### Impact of Clonality

We examined how clonality influences the performance of each method. The results are summarized in Figure 4. Notably, no clear trend emerges between clonality and tree accuracy, likely because none of the models explicitly incorporates clonality into their inference process. As a result, the performance of all methods remains relatively consistent across different clonality levels. NestedBD-Long exhibits particularly low variability, demonstrating robustness in handling both clonal and non-clonal data. In contrast, MEDICC2 and Lazac display higher variability, especially at increased clonality levels, suggesting less stable performance. These findings highlight that while clonality does not directly affect tree accuracy, maintaining stable performance across varying clonality levels is crucial for reliable phylogenetic inference, particularly when clonality is not explicitly modeled.

**Fig. 4.**
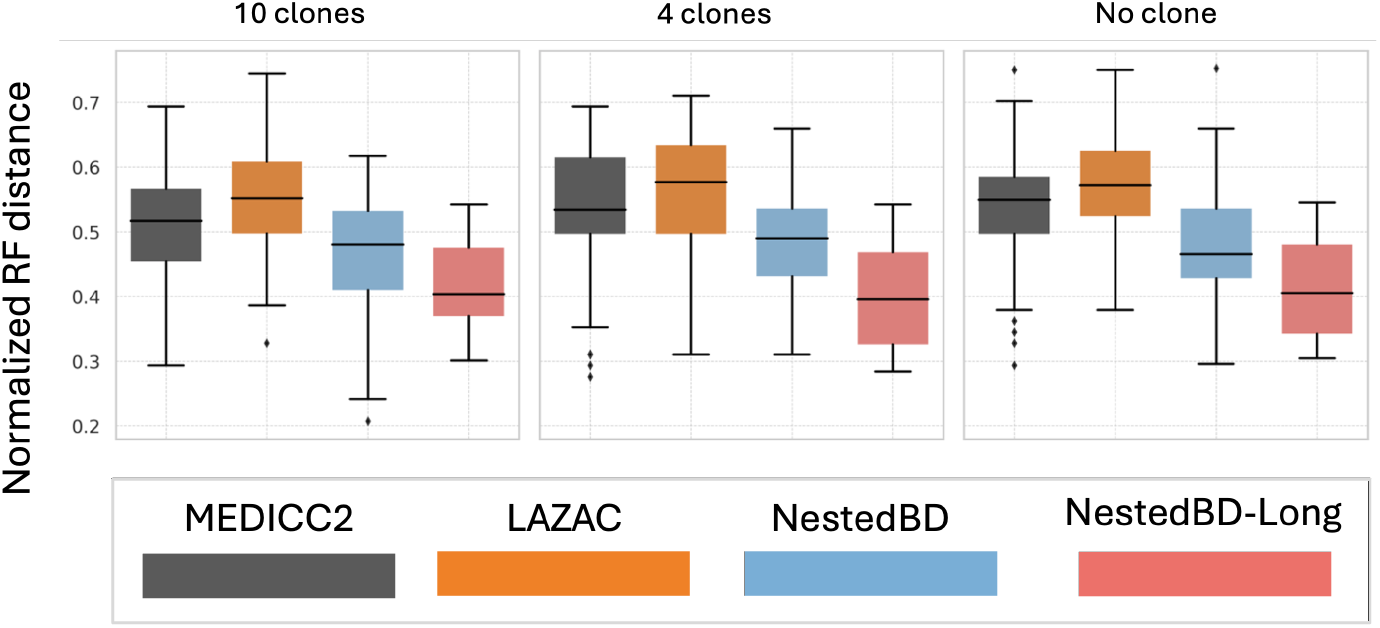
Accuracy of the inferred trees on the simulated data by Clonality. The box plots summarize the distributions of normalized RF distances between the ground truth tree and the inferred trees for each method using 10 replicates.

#### Impact of Error in CNA Profiles

The effect of varying error levels in CNA profiles on tree inference accuracy is summarized in Figure 5. As expected, all methods exhibit increased normalized RF distance with higher error levels, indicating a decline in reconstruction accuracy as data errors increase. NestedBD-Long and NestedBD demonstrate greater robustness, maintaining consistently lower RF distances across different error conditions. In contrast, Lazac shows significant variability, particularly struggling in high-error conditions, suggesting reduced reliability in noisy data. These results indicate that Bayesian methods like NestedBD-Long and NestedBD offer improved stability and resilience to errors, making them more suitable for phylogenetic inference in error-prone single-cell datasets.

**Fig. 5.**
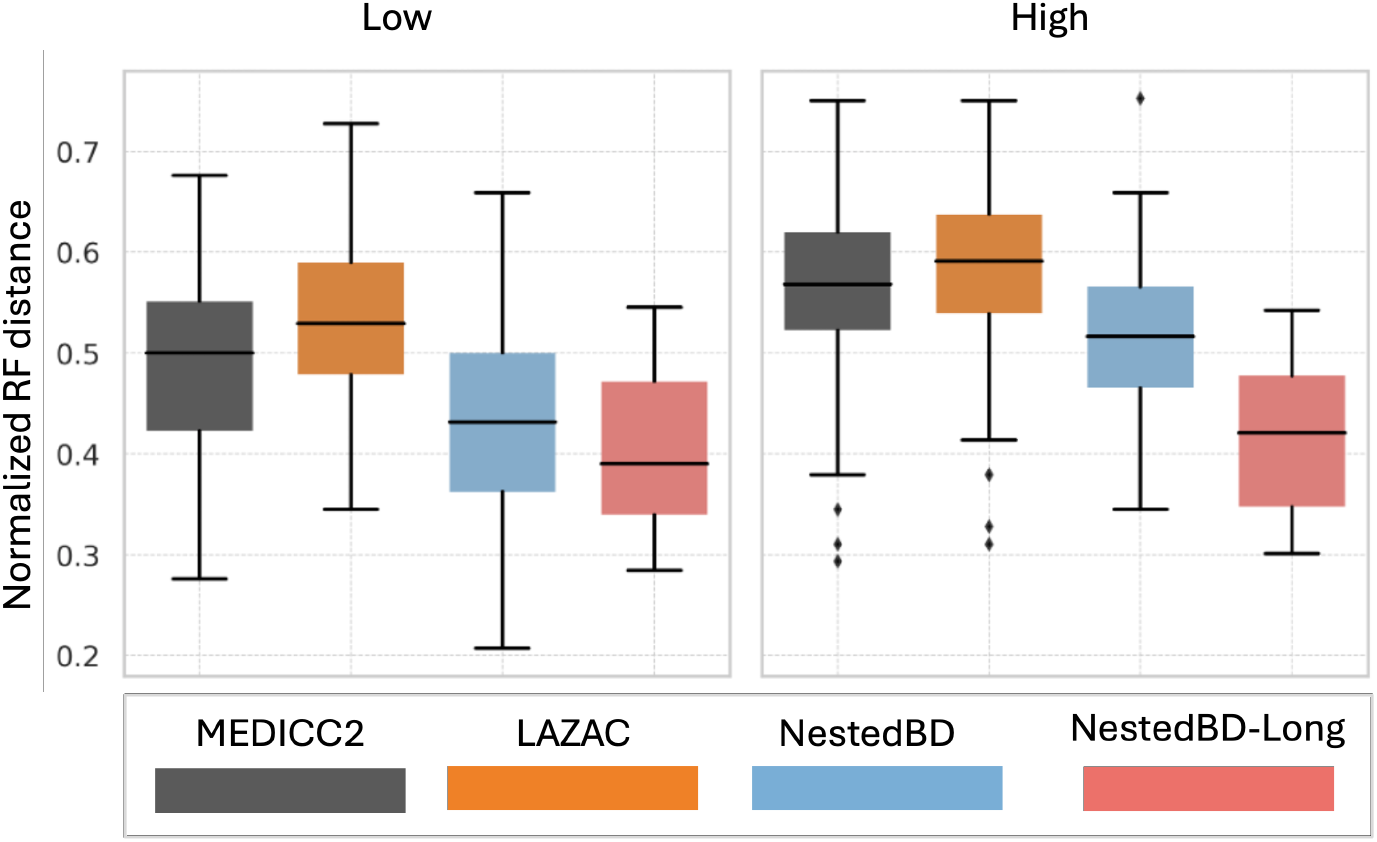
Accuracy of the inferred trees on the simulated data by level of error in CNA profile. The box plots summarize the distributions of normalized RF distances between the ground truth tree and the inferred trees for each method using 10 replicates. ‘High’ and ‘Low’ correspond to the error levels in the simulated CNA profiles (see Section Simulating CNAs).

### 3.2 Performance on Biological Data

We applied all four phylogenetic inference methods to single-cell copy number profile datasets from two TNBC patients, KTN132 and KTN152, obtained from [16]. These datasets contain longitudinal samples from two and three time points, respectively. For each dataset, NestedBD and NestedBD-Long were run for 80 million iterations, with the first 20% of samples discarded as burn-in. The posterior distribution was summarized using the MCC tree, with median node heights. To assess how phylogenetic structure aligns with sampling time points, we computed the pairwise distances between nodes sampled at the same time point and across different time points, normalizing by the largest distance for each method. The results are summarized in Figure 6.

**Fig. 6.**
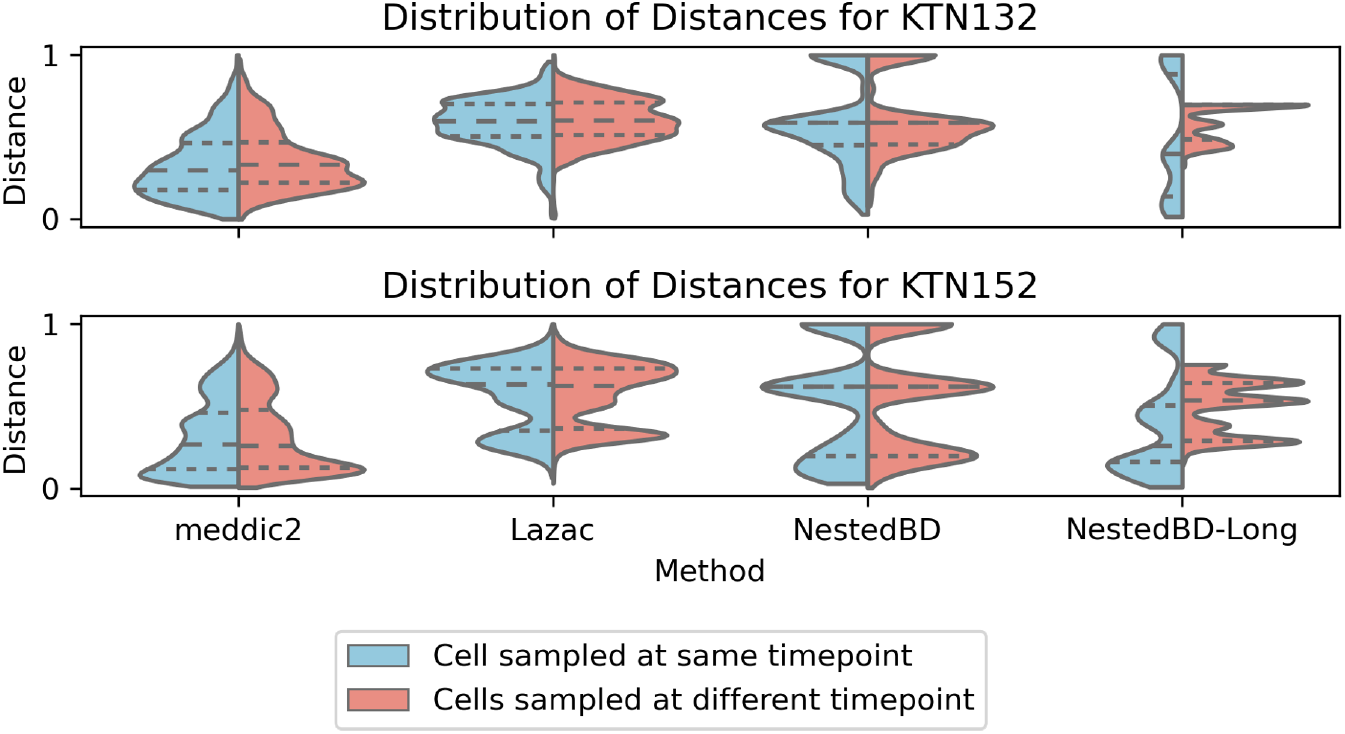
Normalized Pairwise Distance Distributions by Sampling Timepoints. This violin plot illustrates the distribution of normalized pairwise distances between nodes sampled at the same and different time points. The analysis is conducted using four methods — MEDICC2, Lazac, NestedBD, and NestedBD-Long — on singlecell copy number profiles from TNBC patients KTN132 and KTN152 from [16].

For methods that do not explicitly incorporate temporal information, including MEDICC2, Lazac and NestedBD, the distribution of phylogenetic distances between samples from the same and different time points are largely overlapping. It is worth noting that NestedBD inferred a tree with pairwise distances grouped into two or three peaks, which may reflect differences in how it handles rate variation with the birth-death evolutionary model compared to the distancebased method. In contrast, NestedBD-Long, which explicitly models temporal constraints, produces markedly different distributions for pairwise distances between cells sampled at the same time point and those sampled at different time points. This suggests that integrating sampling time allows NestedBD-Long to better capture temporal structure in evolutionary relationships. These results highlight the potential impact of incorporating temporal information when analyzing longitudinal single-cell data.

For each inferred tree, we annotated branches defining major cell clades with cancer-related oncogenes and tumor suppressor genes (TSGs) affected by CNAs, based on data from [26]. The phylogenies inferred using NestedBD-Long for KTN132 and KTN152 are shown in Figure 7 and Figure 8.

**Fig. 7.**
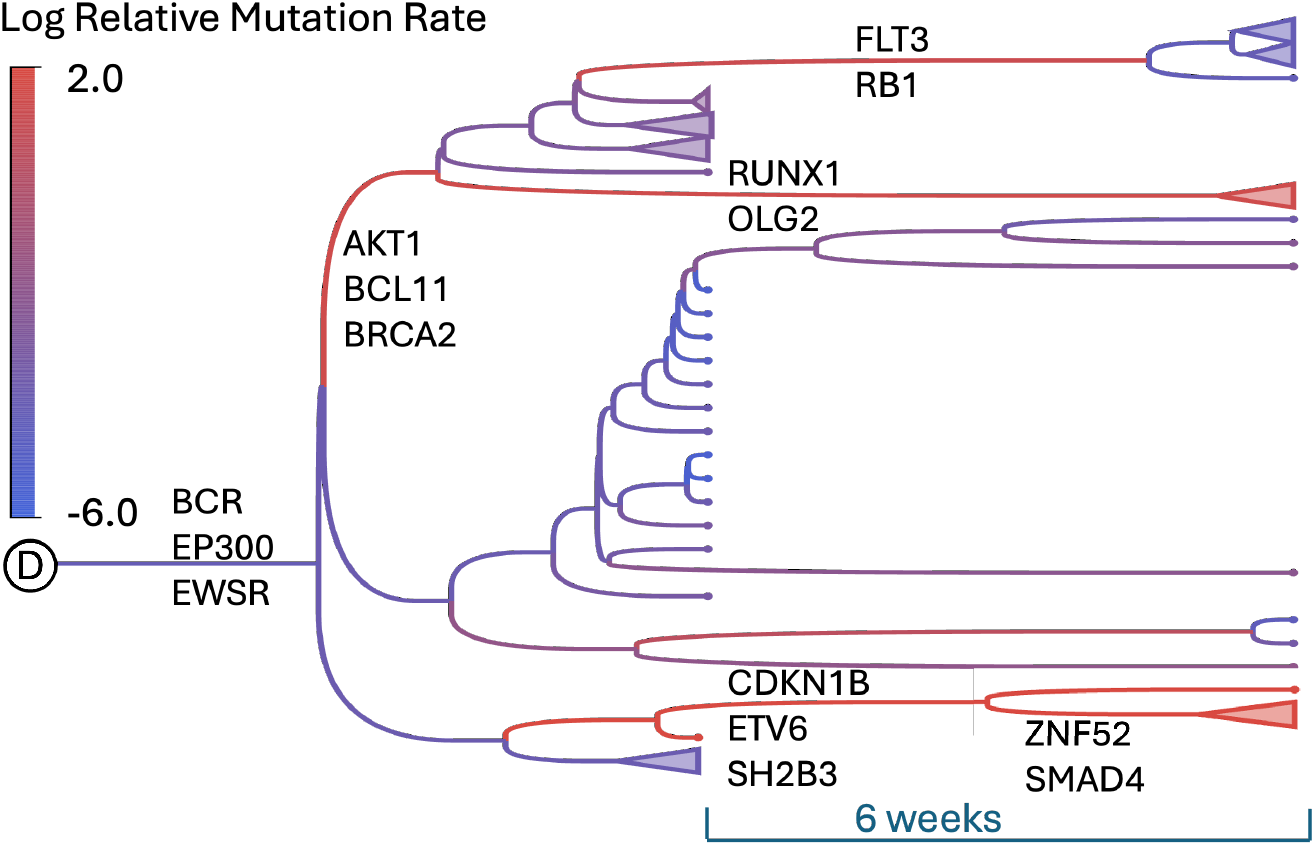
Inferred tree using NestedBD-Long for TNBC patient KTN132. Phylogenetic tree inferred using NestedBD-Long for patient KTN132. Branches defining major cell clades are annotated with TNBC-related oncogenes and tumor suppressor genes (TSGs) impacted by CNAs.

**Fig. 8.**
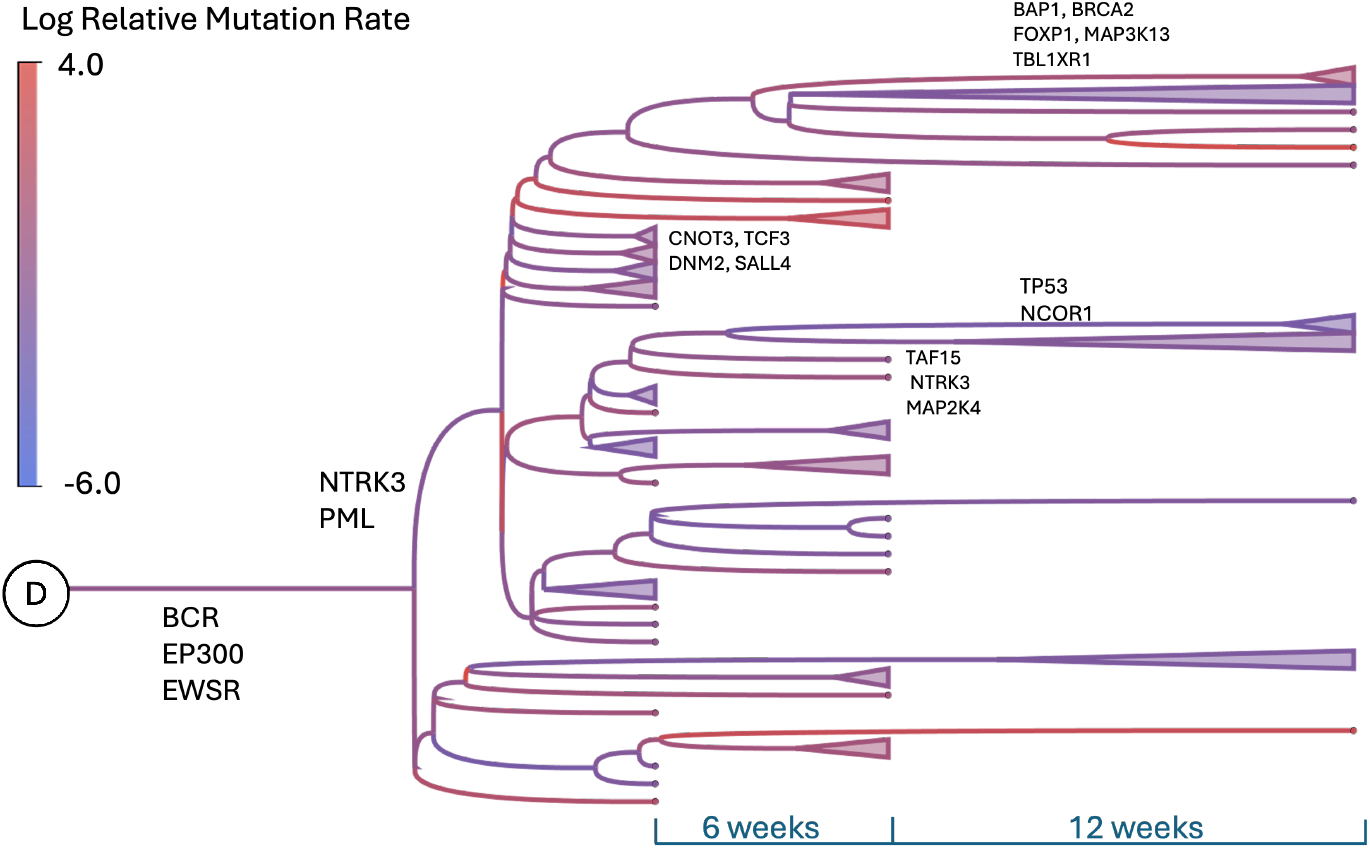
Inferred tree using NestedBD-Long for TNBC patient KTN152. Phylogenetic tree inferred using NestedBD-Long for patient KTN152. Branches defining major cell clades are annotated with TNBC-related oncogenes and tumor suppressor genes (TSGs) impacted by CNAs.

It is worth noting in both figures, branches leading to major clones between sampling points consistently show high mutation rates, indicating rapid clonal evolution and genomic alterations within short intervals. Without longitudinal sampling, scaling mutation rates per unit time would be challenging, risking misinterpretation of clonal expansion dynamics. By integrating multiple time points, NestedBD-Long enables precise temporal reconstruction, distinguishing fast-growing clones from slower-diverging lineages. This is crucial for understanding the timing of key oncogenic events and their role in tumor progression.

For patient KTN132, the inferred tree highlights a subclonal lineage with the RB1 mutation. The high mutation rate on this branch suggests that RB1 loss, known to promote metastasis [12], may have facilitated rapid clonal expansion. We also observed another subclone with the SMAD4 mutation emerging after the first treatment. Its role in drug resistance [29] aligns with the fact that the mutation occurred later in time, likely as a response to treatment. A similar trend is observed in patient KTN152, where branches with key mutations and rapid clonal expansion exhibit increasing mutation rates. Notably, a subclone carrying a TCF3 mutation, implicated in the control of breast cancer growth and initiation[24], shows accelerated mutation accumulation, resembling the pattern seen in KTN132. Additionally, mutations associated with drug resistance including SALL4, FOXP1, and DNM2, emerge at later time points, suggesting a potential response to treatment-induced selective pressures [4,7,5]. This temporal pattern suggests distinct selective pressures acting at different stages of tumor evolution, and reinforces incorporating time information in phylogenetic inference is essential for distinguishing early mutations that drive metastasis from later adaptations to therapeutic pressure.

## 4 Discussion

Longitudinal single-cell sequencing has become an essential tool for tracking tumor evolution and treatment response by capturing dynamic changes in clonal architecture over time. However, existing phylogenetic methods often overlook the importance of explicitly incorporating sampling time, potentially limiting their accuracy in reconstructing evolutionary trajectories. In this study, we introduced NestedBD-Long, which applies the birth-death skyline model to CAN data, integrating temporal sampling to improve evolutionary inference and address limitations in existing methods.

Our evaluations on simulated datasets demonstrate that as the number of temporal sampling points increases, the gap between methods opens up, with NestedBD-Long consistently achieving the highest accuracy. Unlike methods that do not incorporate temporal data, which show increased variability and declining performance, NestedBD-Long maintains stable and precise phylogenetic reconstructions. These results highlight the advantage of explicitly modeling temporal information, particularly in complex evolutionary settings where accurate lineage tracing is critical. On biological datasets, we observed that NestedBD-Long produces distinct distributions of pairwise distances between cells sampled at the same and different time points, demonstrating its ability to capture temporal structure in evolutionary relationships. In contrast, methods that do not account for sampling time exhibit overlapping distance distributions, suggesting a lack of temporal resolution. Furthermore, by following longitudinal constraints, NestedBD-Long can provide more biologically meaningful insights into clonal evolution.

While NestedBD-Long improves temporal resolution in tumor phylogenies, future work could further refine its inference by incorporating additional evolutionary factors such as selection pressures and subclonal interactions. Additionally, extending this framework to infer phylogenies from SNV data or enabling joint inference from both SNV and CNA data could provide a more comprehensive view of tumor evolution. Overall, these findings emphasize the importance of integrating sampling time into phylogenetic reconstruction. By explicitly modeling longitudinal data, NestedBD-Long enhances the accuracy and stability of phylogenetic inference, making it a valuable tool for studying cancer evolution and treatment response.

## Acknowledgments

This research was supported in part by the National Science Foundation, grants IIS-1812822 and IIS-2106837 (L.N.).

